# *TCF7L2* Silencing Reprograms the 4D Nucleome of Colorectal Cancer Cells

**DOI:** 10.1101/2020.05.12.090845

**Authors:** Markus A. Brown, Gabrielle A. Dotson, Scott Ronquist, Georg Emons, Indika Rajapakse, Thomas Ried

**Affiliations:** Genetics Branch, Center for Cancer Research, National Cancer Institute, NIH, Bethesda, MD 20894, USA; Department of Cell Biology and Molecular Genetics, University of Maryland, College Park, MD 20742, USA; Department of Computational Medicine & Bioinformatics, University of Michigan, Ann Arbor, MI 48105, USA; Department of General, Visceral and Pediatric Surgery, University Medical Center, 37075 Göttingen, Germany; Department of Mathematics, University of Michigan, Ann Arbor, MI 48109, USA

## Abstract

Canonical Wnt signaling is crucial for intestinal homeostasis as the major Wnt signaling effector in the intestines, TCF4, is required for stem cell maintenance. The capability of TCF4 to maintain the stem cell phenotype is contingent upon β-catenin, a potent transcriptional activator which interacts with histone acetyltransferases and chromatin remodeling complexes. In colorectal cancer, mutations result in high levels of nuclear β-catenin causing aberrant cell growth. Here, we used RNAi to explore the influence of TCF4 on chromatin structure (Hi-C) and gene expression (RNA sequencing) across a 72-hour time series in colorectal cancer. We found that TCF4 reduction results in a disproportionate upregulation of gene expression genome-wide, including a powerful induction of *SOX2*. Hi-C analysis revealed a general increase in chromatin compaction across the entire time series, though this did not influence gene expression. Analysis of local chromosome organization demonstrated a TAD boundary loss which influenced the expression of a cluster of *CEACAM* genes on chromosome 19. Four-dimensional nucleome (4DN) analysis, which integrates structural (Hi-C) and functional (RNA sequencing) data, identified EMT and E2F as the two most deregulated pathways and *LUM, TMPO*, and *AURKA* as highly influential genes in these networks. Results from gene expression, chromatin structure, and centrality analyses were then integrated to generate a list of candidate transcription factors for reprogramming of colorectal cancer cells to a vulnerable state. The top ranked transcription factor in our analysis was c-JUN, an oncoprotein known to interact with TCF4 and β-catenin.

## Introduction

The Wnt signaling transcription factor, TCF4, is crucial for homeostasis of the mammalian intestine [1]. Loss of TCF4, in either embryonic or adult mice, results in ablation of the proliferative compartment of the intestine [2]. While TCF4 is present throughout the intestine, Wnt signaling is only active at the crypt base where βcat/TCF4 complexes drive a crypt progenitor phenotype [3, 4, 5]. β-catenin is a potent transcriptional activator which influences the surrounding chromatin by recruiting histone acetyltransferases (HATs), chromatin remodeling factors, and RNA polymerase associated factors [6, 7, 8, 9]. Therefore, the binding of a βcat/TCF4 complex significantly enhances the recruitment of the cellular machinery necessary for transcription at Wnt target loci.

In colorectal cancer, mutations in the Wnt signaling pathway, primarily in *APC*, result in high levels of nuclear β-catenin [10, 11]. This results in constitutive Wnt signaling activity and high expression of Wnt target genes, such as *MYC* and *CCND1* [12, 13, 14, 15]. Epigenetic modifications and chromatin remodeling occur as a prerequisite for the expression of Wnt target genes [8, 16]. The high levels of nuclear β-catenin in colorectal cancer likely cause widespread changes in chromatin structure.

It has become evident that understanding chromatin structure lends insight into the regulation of gene expression [17, 18]. Aspects of chromatin structure include the accessibility of genomic loci to the transcriptional machinery as well as the three-dimensional configuration of the chromatin, which may facilitate cooperativity between genomic regions sharing a particular three-dimensional space. Division of the genome into euchromatin and heterochromatin domains, referred to as A and B compartments, respectively, lends insight into the interaction between chromatin structure and function at the chromosome level [17]. Chromosome folding brings distant sites along the linear genome in close spatial proximity, forming topologically associating domains (TADs), which are insulated regions of the genome sharing epigenetic modifications, gene expression, and replication timing [19, 20, 18]. Given the influence of chromatin structure on gene expression, the dissection of a cellular response to a stimulus requires an understanding of both structural and functional dynamics [21, 22].

Integrating chromatin structure and transcriptional dynamics is the basis for the identification of factors to be used in cellular reprogramming [23]. The goal of cellular reprogramming is the guided conversion, typically mediated by transcription factors, of one cell type into another [24, 25, 26]. However, reprogramming also refers to the forced change from one cell state into another, such as converting the cell from a treatment-resistant to a treatment-sensitive state, as this requires the rewiring of genetic circuits and represents an alteration in the response to a stimulus. Unfortunately, the temporal gene expression and chromatin structure data necessary for such an analysis is sparse.

Herein, we explored the dynamical influence of silencing *TCF7L2*, the gene encoding TCF4, on chromatin structure and gene expression across a 72-hour time series in the colorectal cancer cell line, SW480. Silencing of *TCF7L2* not only allows us to investigate the changes which occur as a result of silencing a major transcription factor, it also represents a clinically relevant model as *TCF7L2* expression correlates with resistance to chemoradiotherapy (CRT), a treatment modality in rectal cancer [27, 28, 29]. We were therefore able to track the changes which occurred as cells became increasingly susceptible to CRT. For the time series, an inverse RNAi transfection protocol was designed and optimized, which significantly reduced confounding factors typically found in time series data. A 4DN approach to data analysis was used, which integrated structural (Hi-C) and functional (RNA sequencing) data and allowed us to identify highly influential genes in the most dynamically behaving networks (Fig.1A). We then identified candidate reprogramming factors, to be perturbed alongside *TCF7L2* to further debilitate the colorectal cancer cell.

**Figure 1.**
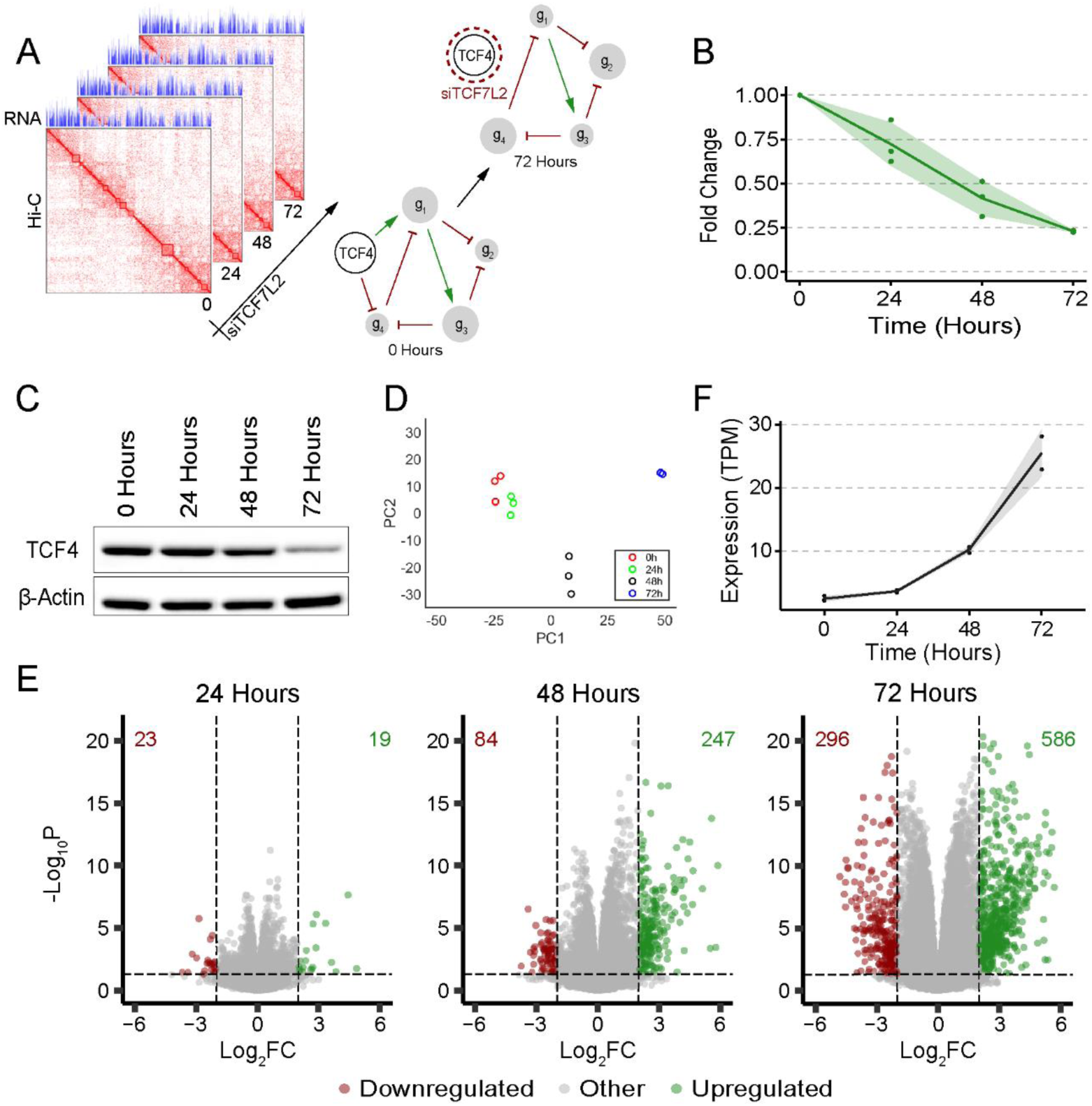
Conceptual Approach and Gene Expression Dynamics Following *TCF7L2* Silencing. (A) Schematic summary of the experimental approach. We sought to explore how silencing *TCF7L2* impacted the colorectal cancer gene network in terms of chromatin structure and gene expression. (B) qPCR demonstrated progressive silencing of *TCF7L2* over time with a 75% reduction in *TCF7L2* transcript levels after 72 hours. Each dot represents a biological replicate, with three replicates plotted. The green line represents the mean, while the green-shaded ribbon represents the standard deviation. (C) Western blot analysis demonstrated a decrease in TCF4 protein abundance over time with β-actin as loading control. (D) PCA of gene expression (TPM) over the time course, with time points differentiated by color. Three replicates are plotted for 0, 24, and 48 hours, while two replicates are plotted for 72 hours. (E) Volcano plots show differential gene expression genome-wide with thresholds set at log_2_FC = 2, and *p* = 0.5 (equivalent to 1.3 on the –log10*P* scale). Genes in red are significantly downregulated and undergo a >4-fold decrease in expression, while those in green are significantly upregulated and experience a >4-fold increase in expression. The number of genes which fall within these regions are shown. (F) Expression profile of *SOX2*, which increases dramatically across the time series. The dots correspond to each biological replicate, the shaded ribbon represents the standard deviation.

## Results

### Global Gene Expression is Upregulated Across the Time Series

The time series data were generated by sequential addition of a small, interfering RNA (siRNA) targeting the 3’ UTR of *TCF7L2* at equally spaced time points (0, 24, 48, and 72 hours) according to a modified siRNA transfection protocol (Fig.S1A, see Methods). The modified protocol was designed to mitigate varying cell cycle distributions, which occur naturally due to varying growth times and are a major confounding factor in extended (>48 hours) time series data (Fig.S1C and D). The modified protocol resulted in ∼70% of the cells in the G_1_ phase of the cell cycle at each time point (Fig.S1B).

To assess the efficacy of siRNA-mediated *TCF7L2* silencing, the quantity of *TCF7L2* transcripts was determined using quantitative PCR (qPCR). Transcript levels decreased progressively across the time points resulting in an ∼2-fold decrease at 72 hours (Fig.1B). The amount of TCF4, the protein product of *TCF7L2*, was determined using Western blot analysis. TCF4 protein levels decreased marginally after 24 hours, ∼35% after 48 hours and ∼70% after 72 hours (Fig.1C). Principal component analysis (PCA) of the RNA sequencing data demonstrated a directed change in the global gene expression program, with biological replicates clustering together (Fig.1D). The number of differentially regulated genes which exhibited both a statistically significant (*p*<0.05) and a 4-fold change in expression increased dramatically across the time series (Fig.1E). However, the balance was positively skewed with nearly twice the number of genes upregulated than downregulated. Considering that SW480 cells have constitutive Wnt signaling activity, and therefore persistent transcription through βcat/TCF4 complexes, the global increase in gene expression following the reduction in TCF4 is unexpected.

### *SOX2* Upregulation is a Result of *TCF7L2* Silencing

To determine the biological relevance of the upregulation of global gene expression, the expression profiles of individual genes with known biological roles were assessed. The expression of the prominent Wnt signaling target genes, *MYC* and *CCND1*, exhibited expression profiles which closely matched that of *TCF7L2* across the time series (Fig.S2)[14, 30, 31]. It has been previously reported that loss of βcat/TCF4-mediated transcription results in the upregulation of genes associated with intestinal differentiation [5]. We confirm this observation as genes associated with differentiation, such as keratin 19 and intestinal alkaline phosphatase, were significantly upregulated following *TCF7L2* silencing (*p*<0.05).

We identified a powerful (log2FC ∼3.35) upregulation of *SOX2*, which began after 24 hours and continued to increase until 72 hours post-transfection (Fig.1F). TCF4 has been previously reported to bind *SOX2* in colorectal cancer [32]. *SOX2* is a transcription factor involved in the regulation of embryonic development, the determination of cell fate, and stem cell maintenance [33]. Interestingly, *SOX2* is one of the four transcription factors required for the reprogramming of adult human fibroblast cells to a pluripotent state [26]. The observation that *TCF7L2* silencing results in the overexpression of *SOX2* suggests that *TCF7L2* may enforce a transcriptional module which potently represses *SOX2* under physiological conditions.

### A/B Compartments are Maintained Over Time

Hi-C was performed to capture genome-wide changes in chromosome structure across the time series to determine the influence of TCF4 on chromatin organization. *TCF7L2* silencing resulted in a significant, genome-wide increase in both the number and strength of pairwise chromatin interactions, as evidenced by matrix subtraction between 0 and 72 hour Hi-C contact maps (Fig.2A). The contact maps demonstrate a higher interaction frequency at 72 hours, while the algebraic difference between the two contact maps reveals that the overwhelming direction of change is an increase in chromatin interaction from 0 to 72 hours. Out of all possible 1Mb-scale chromatin interactions, SW480 cells at 0 hours harbored 74% of those interactions, whereas by 72 hours the proportion of interactions had risen to 87%, demonstrating that the chromatin became more compact over time.

**Figure 2.**
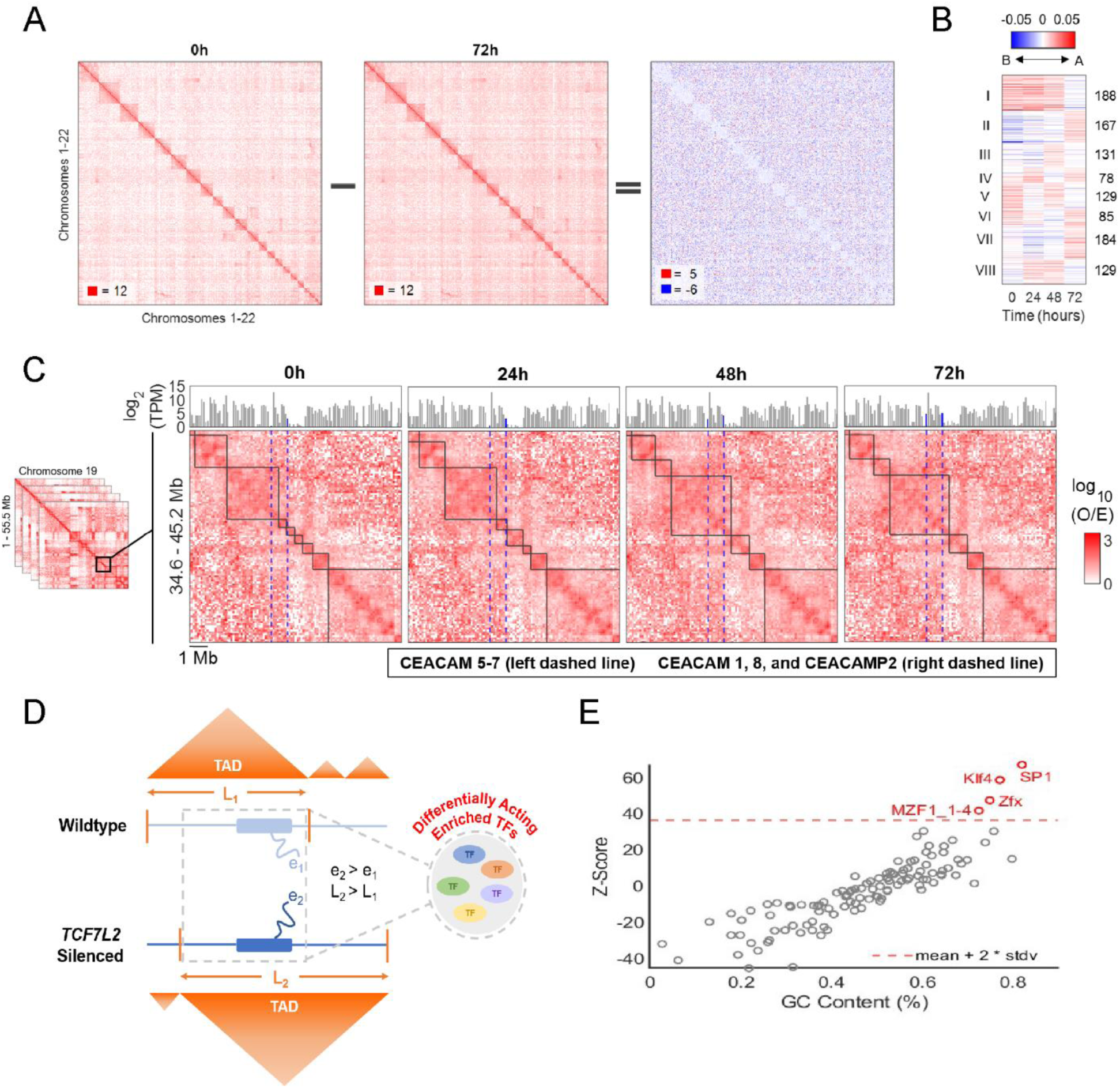
Changes in Global and Local Hi-C Partitioning Over Time (A) Algebraic difference between genomewide Hi-C contact maps at 0 and 72 hours demonstrates an overall increase in contact frequency over time. (B) Global partitioning (A/B compartmentalization) dynamics revealed that 1,091 100-kb genomic bins switched compartments at one or more time points genome-wide. The Fiedler number (see Methods) of each loci are k-means clustered into eight groups (denoted to the left) exhibiting different switching dynamics. The number of genomic bins in each cluster is specified on the right. A 95% change in Fiedler number from one time point to another was set as a threshold to remove inherent noise. (C) Local partitioning (TAD organization) of the region on Chromosome 19 (34.6 - 45.2 Mb) containing two *CEACAM* gene groups. Hi-C contact maps are shown at 100kb resolution with TAD domains denoted by black, solid lines and the *CEACAM* gene groups denoted by blue, dashed lines. The corresponding log2-normalized expression values are highlighted in blue in the panels above. (D) Diagram illustrating coupled chromosome structure and gene expression between two experimental states as well as a possible explanation for this coupling, i.e., cell state-specific transcription factor activity. (E) Transcription factor enrichment analysis for the 64 genes found in the differentially conformed region of chromosome 19 (see 2C) showed enrichment for SP1, KLF4, ZFX, and MZF1 when compared against a background of 24,752 genes.

To further assess changes in genome-wide chromatin organization, the genome was assigned to A/B compartments using a spectral graph theory approach, which calculates the Fiedler number for each 100kb bin (Fig.S3)[34]. The sign of the Fiedler number, positive or negative, indicates within which compartment the bin resides. An A/B compartment switch occurred in ∼4% of the genome at any given point during the time series. K-means clustering identified eight distinct switching patterns in which a specific genomic locus switched compartments unidirectionally (A-to-B or B-to-A) or bidirectionally (a permutation of A-to-B-to-A-to-B) over time (Fig.2B). Approximately 52% of the regions demonstrating an A/B compartment switch underwent a single switch, while the remaining 48% underwent multiple switching events. Despite the switching events, the overall proportion of A/B compartments genome-wide remained relatively unchanged with most loci residing in the same chromatin state across the time series (52% A and 48% B)(Fig.S4).

### TAD Boundary Loss in the *CEACAM* Gene Cluster

Local partitioning of chromatin into topologically associated domains (TADs) provided insight into the coupling of structure and function. Genes occupying the same TADs have been shown to be co-expressed, likely by accessing the same cluster of transcriptional machinery as a result of their spatial proximity [18, 35]. On chromosome 19, an ∼10Mb span of the genome houses a group of *CEACAM* family genes, which are involved in colorectal cancer progression and metastasis [36]. At the initial time point, *CEACAM5, CEACAM6*, and *CEACAM7* were located within the same TAD, while *CEACAM1* and *CEACAM8*, were located in an adjacent, smaller TAD (Fig.2C). Changes in expression of the two *CEACAM* gene clusters after 24 hours were negligible. However, at the 48 hour time point, the boundary separating the two TADs was lost, joining *CEACAM5, CEACAM6*, and *CEACAM7* into a larger TAD with *CEACAM1* and *CEACAM8*. Combination of the TAD domains as well as a significant increase in expression for both *CEACAM* groups occurred seemingly simultaneously at 48 hours. Gene expression for both groups continued to increase following the TAD boundary loss (Fig.2C). The increase in gene expression likely represents increased accessibility to the transcriptional machinery for both gene clusters.

### TAD Boundaries Show Enrichment for the SP1, KLF4, ZFX, and MZF1 Transcription Factors

To identify which transcription factors may be mediating this TAD partitioning loss, a transcription factor enrichment analysis was performed using oPOSSUM-3 (Fig.2D)[37]. Over-representation of transcription factor binding sites in the DNA sequences of the 64 genes occupying the differentially conformed region of chromosome 19 were investigated. The top four identified transcription factors were SP1, KLF4, ZFX, and MZF1 (Fig.2E). The top hit, SP1, is expressed ubiquitously and is involved in chromatin remodeling. KLF4 is involved in embryonic development and binds a GC-rich motif highly similar to SP1. ZFX plays a role in self-renewal of hematopoietic stem cells and MZF1 has been associated in inflammatory bowel disease. Our results indicate that SP1 is the most likely transcription factor mediating this local structural change.

### Coordinated Pathway Responses Suggest a Loss in Cell Cycle Progression and DNA Synthesis Capabilities

To assess the dynamics of TCF4 loss on pathway behavior, fast Gene Set Enrichment Analysis (fGSEA) was performed using the 50 Hallmark gene sets and the log2FC expression values from each time point to generate a normalized enrichment score (NES) [38, 39]. The NESs from fGSEA were compared and the 20 gene sets (10 upregulated and 10 downregulated) with the greatest change by the last time point were plotted (Fig.3A). Strong decreases in expression for both MYC Target gene sets as well as the E2F Targets and G2M Checkpoint gene sets were observed. All four of these gene sets are involved in growth and the regulation of cell cycle progression. The E2F pathway is additionally involved in DNA synthesis, thereby linking DNA synthesis to cell cycle progression [40, 41].

**Figure 3.**
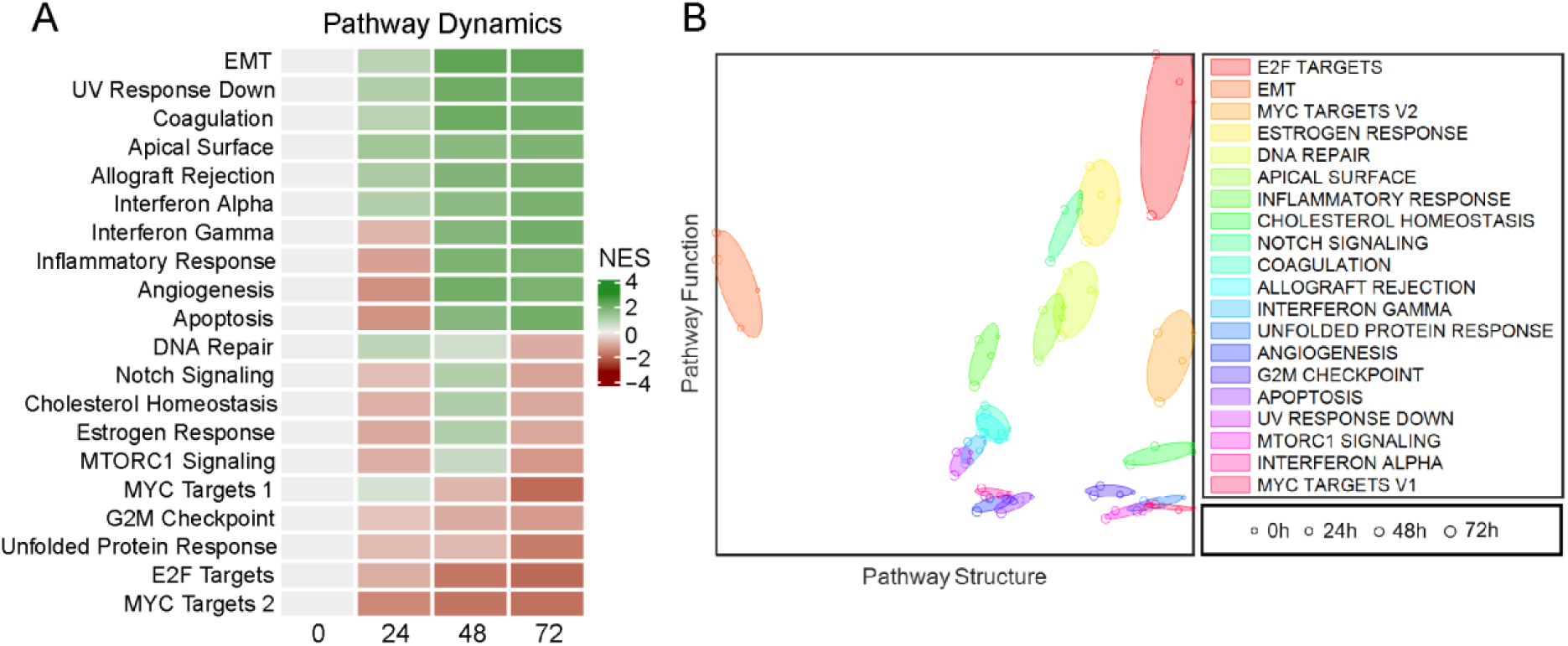
Pathway Level Gene Expression and Structure-Function Analyses. (A) Fast Gene Set Enrichment Analysis (fGSEA) was performed using the Hallmark Gene Sets and log_2_FC for each time point to generate a normalized enrichment score (NES). The NESs of the ten most up- and downregulated pathways were then plotted as a heatmap to show pathway level gene expression over time. The NES at the initial time point is set to 0. (B) Structure-function analysis of the pathways shown in (A) compares Pathway Structure (Frobenius norm of chromatin interaction frequencies) and Pathway Function (Frobenius norm of gene expression). Each pathway creates an ellipse bounded by the structure-function values at each time point. Increasing time points are shown as circles of increasing diameter, with color denoting the pathway. The E2F, EMT, and MYC Targets V2 are the most dynamically changing pathways in terms of their structure-function relationship.

Interestingly, the Notch, Cholesterol homeostasis, Estrogen Response, and MTORC1 signaling pathways all demonstrated a similar NES profile over time, suggesting co-regulation. Several articles identify interplay between these pathways, supporting this hypothesis [42, 43, 44]. The Interferon Gamma, Inflammatory Response, Angiogenesis, and Apoptosis pathways also demonstrate a coordinated network of upregulated pathways, which appears to regulate a wound repair axis in colorectal cancer. The most dynamically upregulated gene set was EMT, which suggests that invasion and migration capabilities are increased upon silencing of *TCF7L2*.

### Pathway Level Structure-Function Relationships

We then sought to compare structure-function relationships in these pathways, utilizing a metric called the Frobenius norm, which describes the space of a vector (gene expression) or matrix (chromatin interaction frequencies) [45]. The norm of pathway specific RNA-seq and Hi-C data for each time point was computed by normalizing for the size of the pathway. Min-max normalization was then performed to gauge the correlation between structure and function for any given pathway as well as to make direct comparisons between multiple pathways. The normalized values were then projected onto a phase portrait for visualization (Fig.3B). Pathways are represented as ellipses demarcated by the coordinates of the four time points. Pathways which fall along the diagonal, including the E2F Targets, Estrogen Response, and DNA Repair gene sets, demonstrate a coupling between structure and function, indicating that chromatin organization and RNA transcription are functioning in tandem. Pathways in the lower right quadrant indicate a weaker correlation between structure and function. These pathways demonstrate a chromatin structure which is primed for higher expression, yet currently do not meet the expression levels typically associated with that chromatin structure. The EMT pathway, far left, demonstrates a large change in function yet a marginal change in structure. This indicates that the chromatin was in an accessible conformation for gene transcription prior to the silencing of *TCF7L2*, which likely facilitated its powerful induction.

### EMT and E2F Signaling Genes are Highly Connected in SW480 Cells

The two most dynamic gene sets, EMT and E2F, were then probed to gauge the connection between these networks in three-dimensional space. A Hi-C-derived, synthetic 5C contact map was generated to highlight the inter-loci interaction frequencies among EMT and E2F signaling genes. The contact map is comprised of 1Mb bins containing genes in either of the pathways (see Methods). Compared to a synthetic 5C contact map composed of randomly sampled, gene-containing genomic bins, we observe that the EMT and E2F signaling genes contact map is denser, suggesting that these genes are more closely interacting than other genes in the genome (Fig.4A). Permutation testing reveals that these genes are indeed more inter-connected than genes outside either network (*p* = 0.001), with a Fiedler number that surpasses those of 1,000 sets of randomly sampled genes (Fig.S5A,B).

**Figure 4.**
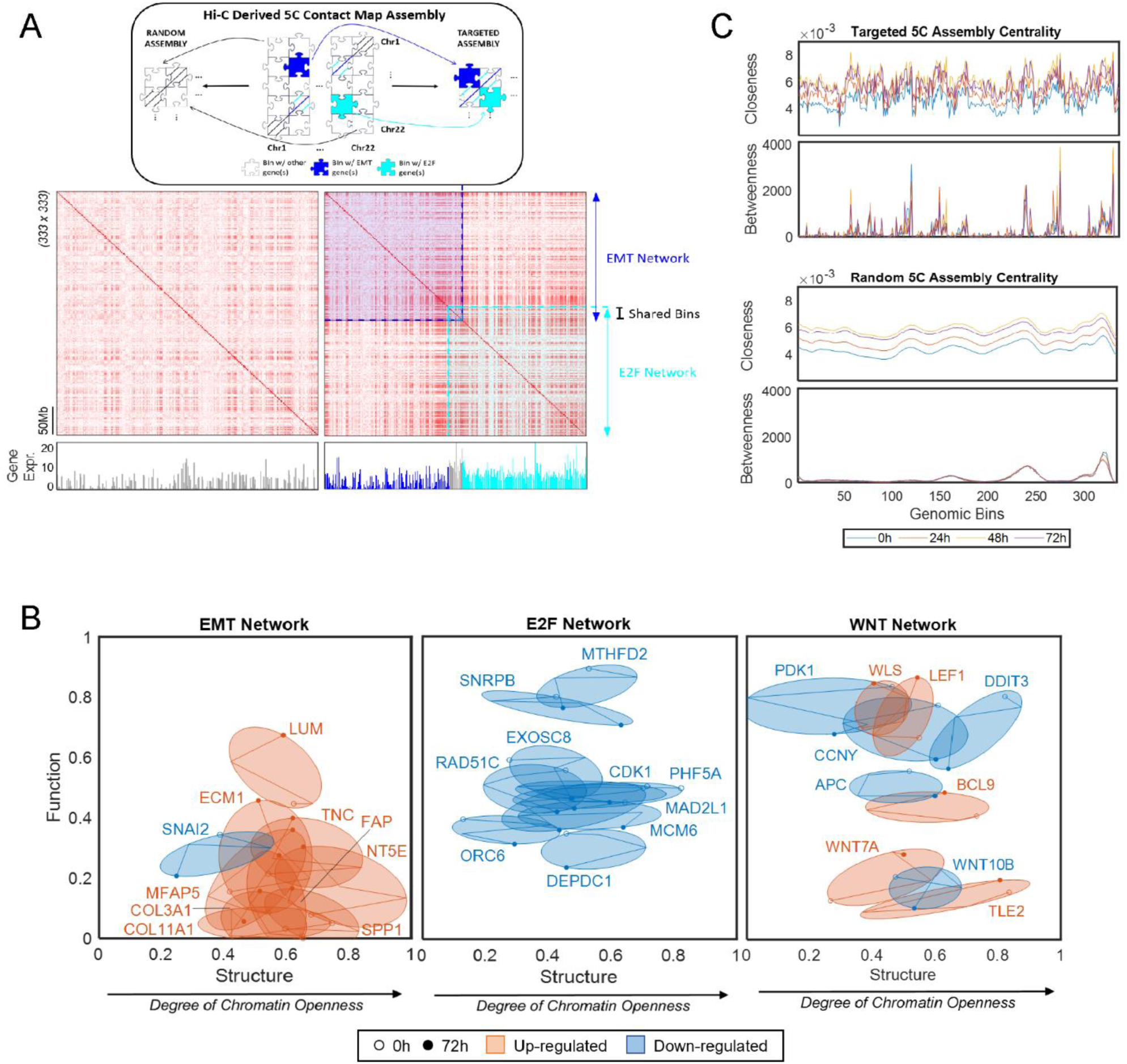
Centrality Analyses and Gene Level Structure-Function Relationships. (A) Gene network-level synthetic 5C contact maps were generated by extracting bins corresponding to genes in the EMT and E2F signaling networks from a 1Mb adjacency matrix and stitched together (top) to create a synthetic contact map (right). Synthetic 5C contact map for randomly sampled gene bins (left) shown for comparison. (B) Structure versus function phase portrait for the EMT, E2F, and WNT signaling pathway genes. Min-Max normalized Von Neumann Entropy (see Methods) and gene expression represent structure and function, respectively. The ten most dynamic genes (those whose fitted ellipse have the largest areas) in each network are shown. Genes that decrease in expression over time are labelled in blue and genes that increase in expression are labelled in orange. Phase portraits reveal the extent to which chromatin structure and gene expression are coupled. (C) Closeness centrality increases over time, indicating that the network becomes increasingly compact. An increase in betweenness centrality is also observed over time which reaches a maximum at 48 hours. Peaks in betweenness centrality plot identify bins (genes) which significantly regulate the network. The Random 5C Assembly demonstrates a progressive increase in closeness centrality, reflective of the general increased contact frequencies over time.

To understand the evolving behavior of individual genes in the EMT, E2F, and Wnt signaling networks, a gene-level, structure-function analysis was performed. Each gene was examined at 5kb resolution and was assigned a 35kb-length window – 5kb overlapping the transcription start site and 15kb flanking either side of that 5kb region – creating a seven by seven sub-matrix for each gene. Several network-based approaches have previously been used to characterize behaviors in dynamically changing genomes [46, 47, 48]. We apply one such approach - a derivative of Von Neumann Entropy (VNE) - to measure local chromatin organization of individual gene regions [49]. Higher VNE values indicate that the number of conformations available to the gene and its immediate neighborhood are higher, indicating that chromatin is more accessible. We compute VNE on the EMT, E2F, and Wnt signaling gene sub-matrices and plot them as a function of gene expression, creating a phase portrait (Fig.4B). This shows the gene-level trajectory of chromatin accessibility and expression for the ten most dynamic genes, as determined by the area of their ellipse. Overlap between the gene ellipses demonstrates a homogeneous concerted pathway response in the EMT and E2F gene sets, indicating that the genes, in their respective pathways, are in a similar chromatin environment and demonstrate coordinated changes in expression. The WNT gene network demonstrates a greater diversity in its response, indicative of varying chromatin environments and diverse transcriptional regulation at the individual gene loci. Genes with different expression patterns, yet whose phase portraits still overlap, are indicative of regulation at the transcriptional level (such as transcription factor binding), rather than at the chromatin level. For example, *CCNY* and *LEF1* are present in similar chromatin environments, as indicated by their overlapping phase portraits, yet show opposite funtional patterns, indicating that their regulation occurs at the transcriptional level. Overall, we observe that the fitted ellipses change dynamically in terms of structure and function, confirming that *TCF7L2* silencing influences chromatin structure.

### Network Interactions are Hardwired and Overall Proximity Increases Over Time

To further explore chromatin topology, we performed network centrality analysis on inter-loci interaction data from our synthetic 5C contact maps. Network centrality describes the connectedness of a network, which yields insight into the path by which information is relayed through the network [50, 51, 47]. We calculated four measures of centrality: betweenness, closeness, degree, and eigenvector (see Methods). Betweenness centrality identifies nodes which lie along the shortest path between other nodes, closeness centrality is determined by the average farness to all other nodes, degree centrality captures the number of edges connected to a node, and eigenvector centrality determines the influence of a node based on the influence of the neighboring nodes.

Roughly 57% of the EMT and E2F network genes underwent a significant change in betweenness centrality, the majority of which experienced a decrease in their betweenness centrality score (Fig.4C). *LUM, TMPO*, and *AURKA* had the highest betweenness centrality scores at any given time point, suggesting that they are highly influential in their respective networks. Since silencing of *TCF7L2* resulted in decreased betweenness centrality scores for these genes, they represent the most likely nodes by which TCF4 may influence the EMT and E2F networks.

Closeness centrality scores provided insight into the evolving proximity of the network genes. We observe that closeness centrality increased with time for all loci with either of the latter two time points (48 and 72 hours) having the highest scores (Fig.4C). This accompanies the previous observation that pairwise chromatin interactions increase over time, i.e., chromatin organization is becoming denser. Since a higher closeness centrality score indicates that a node is relatively closer to all other nodes in the network, this trend suggests that the genes become more organized with time. We observe that the eigenvector and degree centralities of our network genes fluctuate very little with time - meaning the same gene interactions with a similar magnitude of connectivity occur at each time point (Fig.S5C).

### 4DN Analysis Provides Insight into TF-Driven Controllability of CRC Cells

Our findings demonstrate that silencing *TCF7L2* is sufficient to mediate structural and functional changes that influence the behavior of colorectal cancer cells. We suspect that there are a number of reprogramming factors that, when silenced alongside *TCF7L2*, could further destabilize the colorectal cancer cell. Genes that have significant nodal influence represent ideal candidate reprogramming factors, as a result of their ability to destabilize and promote wide-spread changes in the network. Betweenness centrality is one measure of nodal influence and is particularly characteristic of cross-talk control. We find that in the colorectal cancer network, 187 1Mb bins containing 215 genes from the EMT and E2F signaling gene networks harbored a significant change in betweenness centrality from 0 to 72 hours. These genes were ranked according to their magnitude of change in betweenness centrality, with higher magnitudes corresponding to a higher ranking. The genes were then ranked separately by their percent change in gene expression and Frobenius norm between 0 and 72 hours to incorporate aspects of gene function and structure. The average of the three rankings for each gene was computed and the genes ordered accordingly, with lower rankings indicating a gene with the most controllability potential. Considering that transcription factors regulate a network of genes, we summarize the rankings and dynamics for the most likely transcription factor-encoding genes from our candidate gene list in Table 1. The highest ranked candidate was c-JUN, an oncogenic subunit of the AP-1 transcription factor, whose activity is augmented in many cancer types, and has been shown to interact with TCF4 and β-catenin to form a ternary, transcriptionally-competent complex [52].

**Table 1.**
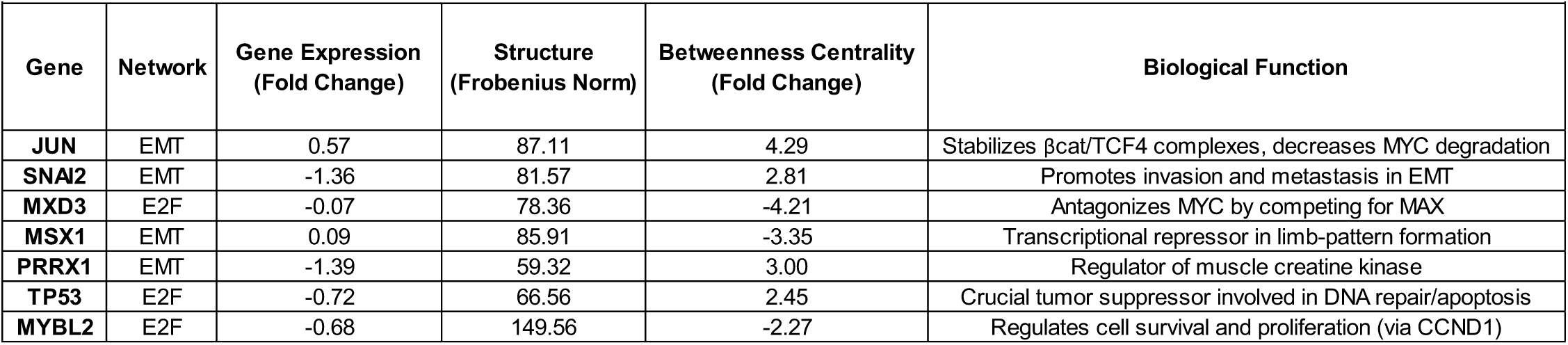
Ranked Candidate Reprogramming Factors. A total of 215 genes from the EMT and E2F signaling networks were ranked according to the magnitude of their change in gene expression, frobenius norm, and betweenness centrality, between the 0 and 72 hour time points, irrespective of direction. The list of candidate factors was then filtered by removing genes which are not well characterized as transcription factors as well as those whose motif binding information was unavailable.

## Discussion

Colorectal cancers exhibit constitutively active Wnt signaling, largely due to mutations in *APC*[10]. The overexpression of Wnt target genes is driven by βcat/TCF4 complexes, as TCF4 is the major Wnt signaling transcription factor in colorectal cancer [13]. Due to the central role of Wnt signaling in colorectal tumorigenesis as well as the capability of β-catenin to modulate the chromatin environment, we explored the influence of TCF4 on chromatin structure and gene expression dynamics in the SW480 colorectal cancer cell line.

*TCF7L2* silencing resulted in a progressive deregulation of the transcriptome with the most dynamic changes occurring at the last time point. Surprisingly, nearly twice the number of genes were upregulated upon *TCF7L2* silencing. While difficult to distinguish between direct and indirect effects, these results hint at a repressive role for TCF4. Indeed, naturally occurring dominant-negative variants of TCF4, have been described [53, 54]. The gene *SOX2* demonstrated a powerful increase in expression which occurred progressively over the time series. Interactions between canonical Wnt and *SOX2* have been described in various cell types. In lung epithelia, an antagonizing role between canonical Wnt signaling and *SOX2* has been described in which activation of Wnt signaling led to the loss of *SOX2* and interfered with the development of the bronchiolar epithelium [55]. In gastric tumorigenesis, *SOX2* functioned as a tumor suppressor which antagonized Wnt-driven adenomas [56]. These two examples demonstrate a mutual, antagonizing relationship between *SOX2* and canonical Wnt signaling, which influences the development of normal tissue (lung) or adenomas (stomach). Here, we observe an antagonizing role between TCF4 and *SOX2* in colorectal cancer. Taken together, these results suggest that TCF4 maintains a transcriptional module for repressing *SOX2*, which may be necessary for generating and/or maintaining the intestinal lineage.

Time series Hi-C experiments demonstrated that TCF4 influences chromatin organization both globally and locally. Silencing of *TCF7L2* led to an increase in pairwise chromatin interactions, demonstrating an overall compaction of the chromatin across the genome. This may be due to a decrease in βcat/TCF4 binding, which would lead to decreased β-catenin-mediated chromatin acetylation. The high levels of nuclear β-catenin present in colorectal cancer may be sufficient for mediating a genome-scale chromatin effect. However, the compaction did not result in a decrease in gene expression.

Approximately 4% of the genome underwent an A/B compartment switch at any given time point. The silencing of a single transcription factor leading to compartment switching in ∼4% of the genome is comparable to overexpressing four transcription factors and influencing ∼20% of the genome [57]. Despite changes in A/B compartment switching, the total number of bins which resided in the A/B compartments remained stable over time (52% A, 48% B). This supports the existence of a genome-wide ‘balance’ of open and closed chromatin domains. However, the A/B compartment balance in normal cells of the mouse is different, at 40% A and 60% B [57]. This may be a reflection of the difference in species or could be an indication that, since cancer cells have enlarged nuclei, they may be able to support a larger percentage of the genome in the A compartment, and therefore support a more diverse transcriptome. However, an increased nuclear volume could merely reflect the increased DNA content of cancer cells.

Local shifts in chromosome partitioning revealed a TAD boundary loss on chromosome 19, which was coordinated with an increase in gene expression in the *CEACAM* family genes. The most dramatic change in gene expression occurred concomitantly as the TAD boundary loss, suggesting a causal link between the two. Following the TAD domain coalescence, the expression of both gene groups continued to increase, likely by accessing the same transcriptional machinery. Given the current understanding of structure-function relationships, we interpret the dynamics of the event thus: *TCF7L2* silencing results in the decreased activity of a factor, likely SP1, which results in loss of the TAD boundary. Boundary loss increases the probability that the two *CEACAM* groups will come in close contact, thereby allowing them to coordinate recruitment of the transcriptional machinery.

From fGSEA, we observed downregulation of the MYC, E2F, and the G2/M gene sets, all which mediate growth and cell cycle progression. In addition to regulating growth, genes in the E2F pathway are also responsible for DNA synthesis [40, 41]. Additionally, despite showing the greatest decrease in pathway gene expression, E2F genes are still expressed to a higher degree than genes in EMT, the most upregulated gene set (Fig.4A - Gene Expression). Previous evidence has demonstrated that TCF4 expression corresponds with resistance to CRT and that loss of TCF4 increases sensitivity [27, 28, 58]. Furthermore, it was been suggested that the mutation rate in intestinal stem cells is relatively equal to the mutation rate in liver stem cells, despite the higher proliferative rate of the former [59, 60]. We hypothesize that high levels of DNA synthesis/DNA proof-reading activity, mediated by E2F, could maintain DNA sequence integrity in the face of continued stem cell proliferation and, in colorectal cancers, this same DNA program would defend against CRT. Loss of TCF4 results in cell cycle arrest, ensuring that cells with hampered DNA repair capabilities do not propagate in the stem cell crypt.

Investigation of structure and function relationships in the top 20 pathways demonstrated that ∼50% of the pathways undergo corresponding changes in pathway structure and function, indicating that the chromatin environment of the genes comprising the pathway is representative of their expression. The EMT pathway shows marginal changes in structure yet demonstrates the largest upregulation in function, indicating that the chromatin conformation for the genes in EMT were primed for expression at the initial time point. This implies that colorectal cancers can activate EMT more rapidly than other pathways, based on the pre-existing conformation of their genome. Closer examination of the EMT and E2F genes demonstrated preferential interaction between one another in three-dimensional space. Considering the negative correlation between these pathways, their association in three-dimensional space is antagonizing and suggests that dynamic behavior in one may be responsible for dynamic behavior in the other.

Centrality analysis allowed us to determine the most influential genes in their respective networks. The genes with the highest betweenness centrality were *LUM, TMPO*, and *AURKA*, identifying these genes as the most useful targets for propagating a stimulus through their respective pathways (EMT or E2F). We subsequently identified a list of factors which can be used in concert with loss of TCF4 to profoundly perturb the colorectal cancer cell. The top ranked candidate reprogramming factor was c-JUN, an oncoprotein which forms part of the AP-1 transcription factor. Previous work has shown that c-JUN forms a complex with TCF4 and β-catenin, which stabilizes β-catenin, and drives expression of the *JUN* promoter, initiating a self-stimulating feedback loop [61, 62]. Abrogation of the c-JUN, TCF4, β-catenin complex led to reduced tumor number and size, as well as prolonged lifespan [52]. High levels of c-JUN and Wnt signaling activity resulted in increased resistance to cisplatin whereas abrogation of c-JUN or Wnt signaling activity increased the sensitivity of ovarian cancer cells to treatment.

In conclusion, we find that *TCF7L2* influences both structure and function in the SW480 colorectal cancer cell line. We find evidence supporting the role of *TCF7L2* in promoting cell growth and DNA synthesis/DNA repair in colorectal cancer due to the decreased expression of the MYC, E2F, and G2M gene sets upon TCF4 reduction. When TCF4 is present, chromatin resides in a more open conformation genome-wide, which is likely mediated by βcat/TCF4 complexes. TCF4 is capable of inducing A/B compartment switching, though is unable to perturb the A/B compartment balance. At the local chromatin level, the expression of similarly regulated genes at a TAD boundary was greatly increased upon TAD boundary loss. The two most dynamic pathways following *TCF7L2* silencing – EMT and E2F – interact in three-dimensional space and the most dynamic pathway genes inhabit overlapping structure-function space. The most influential genes in the network were identified using measures of centrality. Incorporation of chromatin structure, gene expression, and centrality data allowed us to generate a list of transcription factors which would be most effective at perturbing the colorectal cancer cell in conjunction with TCF4. Our top ranked transcription factor, c-JUN, is known to bind TCF4 and β-catenin, thereby confirming the validity of network connectivity analysis in identifying relevant biological interactions.

## Methods

### Cell Culture

The colorectal adenocarcinoma cell line, SW480 (CCL-228) was obtained from the American Type Culture Collection (ATCC). Cells were grown in RPMI-1640 growth medium (Gibco; 11875093), supplemented to 10% FBS (Gibco; 10082147) and 2mM L-glutamine (Gibco; 25030149). Cells were incubated at 37C in 5% CO_2_. No antibiotics were used. Proper cell line identity was confirmed using STR profiling and the absence of mycoplasma contamination was confirmed routinely using PCR (Genelantis; MY01050).

### siRNA Inverted Transfections

Cells were seeded in a 6-well tissue culture plate (Falcon; 353046) at a density of 150,000 cells per well. The wells contained 2mL of RPMI-1640 growth medium. Silencer Select siRNAs (Invitrogen), Negative Control No.1 (4390843) and siTCF7L2-s13880 (4392420), were delivered to the cells using the lipofectamine RNAiMAX reagent (Invitrogen; 13778150), according to the manufacturer’s instructions. This resulted in the addition of 7.5µL of RNAiMAX and 25pmol of siRNA per well of the 6-well plate. The siRNA:lipofectamine complexes were not removed. The time series was constructed thus: 0 and 72 hour time points were transfected with siNEG and siTCF7L2, respectively, 24 hours after seeding. The 48 hour time point was transfected, with siTCF7L2, 24 hours after the 0 and 72 hour time points. The 24 hour time point was transfected, with siTCF7L2, 24 hours after the 48 hour time point. The cells were harvested 24 hours later, resulting in exposure to siTCF7L2 for 72, 48, and 24 hours. The 0 hour time point spent 0 hours in siTCF7L2 and serves as a transfection control for the siTCF7L2 72 hour time point.

### Quantitative PCR

RNA was extracted from at least three independent inverted transfections using the RNeasy Mini Kit and on-column DNAse treatment (Qiagen; 74104, 79254), according to the manufacturer’s instructions. RNA concentration was determined using a NanoDrop 1000 spectrophotometer (Thermo Scientific). Synthesis of cDNA was performed using the Verso cDNA Synthesis kit (Thermo Scientific; AB1453B), based on the manufacturer’s instructions. The optional RT Enhancer was used and equal amounts of both the Anchored oligo dT and Random Hexamers were added. Quantitative PCR was performed for *TCF7L2* and *YWHAZ* on an ABI PRISM 7000 sequence detection system (Applied Biosystems) using Power SYBR Green PCR Master Mix (Applied Biosystems; 4367659), according to the manufacturer’s instructions. Primer sequences can be found in the supplement. The thermocycling protocol consisted of a 10 minute hold at 95C followed by 40 two-step cycles of 15 seconds at 95C and 1 minute at 60C. Fold change was determined using the *YWHAZ* reference gene and 2^-ΔΔCT^.

### Western Blot

Nuclear and cytoplasmic proteins were extracted using the NE-PER Nuclear and Cytoplasmic Extraction Kit (Thermo Scientific; 78833), according to the manufacturer’s instructions. Protein was prepared for quantification using the Pierce BCA Protein Assay Kit (Thermo Scientific; 23225), according to the manufacturer’s instructions, and quantified on a SpectraMAX M2e microplate reader (Molecular Devices). Protein samples (20µg) were electrophoresed using a 4-12% Bis-Tris gel (Invitrogen; NP0322BOX) in a XCell Sure-Lock Mini-Cell (Invitrogen; EI0002), transferred to a PVDF membrane, and incubated with antibodies targeting TCF4 or β-actin (Abcam; 76151 and CST; 4970S, respectively). An HRP-secondary antibody (CST; 7074S) was applied and detected using Pierce ECL Western Blotting Substrate (Thermo Scientific; 32209). Blots were imaged in an Azure c600 Gel Imaging Station (Azure Biosystems).

### RNA Sequencing

RNA was extracted from at least three independent inverted transfections using the RNeasy Mini Kit and on-column DNAse treatment (Qiagen; 74104, 79254), according to the manufacturer’s instructions. RNA concentration was determined using a NanoDrop 1000 spectrophotometer (Thermo Scientific). RNA integrity was determined using the 4200 TapeStation (Agilent). RNA samples with an RNA integrity number of 9 or higher were used for sequencing. Ribosomal RNA (rRNA) was removed using biotinylated, target-specific oligos combined with Ribo-Zero rRNA removal beads according to the Illumina total RNA sequencing protocol. The RNA was fragmented and the cleaved RNA fragments were copied into first strand cDNA using reverse transcriptase and random primers, followed by second strand cDNA synthesis using DNA Polymerase I and RNase H. The resulting double-strand cDNA was used as the input to a standard Illumina library prep with end-repair, adapter ligation and PCR. The final purified product was quantitated by qPCR. Samples were sequenced on a HiSeq2500 using Illumina TruSeq v4 chemistry with 125 bp paired-end. Reads were aligned to reference genome hg19 using STAR. Gene expression was quantified using RSEM. Alignment parameters were set based on RSEM default parameters, “rsem-calculate-expression” “–star”. Transcripts per million (TPM) was used to quantify gene expression.

### Hi-C

The *in situ* Hi-C protocols from Rao et al. were adapted with slight modifications. For each Hi-C library, approximate 3 x 10^6^ cells were incubated in 250µl of ice-cold Hi-C lysis buffer (10mM Tris–HCl pH 8.0, 10mM NaCl, 0.2% Igepal CA630) with 50µl of protease inhibitors on ice for 30 minutes and washed with 250 µl lysis buffer. The nuclei were pelleted by centrifugation at 2500g for 5 minutes at 4C, re-suspended in 50µl of 0.5% sodium dodecyl sulfate (SDS) and incubated at 62C for 10 minutes. Afterwards, 145µl of water and 25µl of 10% Triton X-100 were added and incubated at 37C for 15 minutes.

Chromatin was digested with 200 units of MboI (NEB) overnight at 37C with rotation. Chromatin end overhangs were filled in and marked with biotin-14-dATP (Thermo Fisher Scientific) by adding the following components to the reaction: 37.5µl of 0.4mM biotin-14-dATP (Life Technologies), 1.5µl of 10mM dCTP, 1.5µl of 10mM dGTP, 1.5µl of 10mM dTTP, and 8µl of 5U/µl DNA Polymerase I, Large (Klenow) Fragment (NEB). The marked chromatin ends were ligated by adding 900µl of ligation master mix consisting of 663µl of water, 120µl of 10X NEB T4 DNA ligase buffer (NEB), 100µl of 10% Triton X-100, 12µl of 10mg/ml BSA, 5µl of 400U/µl T4 DNA Ligase (NEB), and incubated at room temperature for 4 hours.

Chromatin reverse crosslinking was performed by adding 50µl of 20mg/ml proteinase K (NEB) and 120µl of 10% SDS and incubated at 55C for 30 minutes, adding 130µl of 5M sodium chloride and incubated at 68C overnight. DNA was precipitated with ethanol, washed with 70% ethanol, and dissolved in 105µl of 10 mM Tris–HCl, pH 8. DNA was sheared on a Covaris S2 sonicator. Biotinylated DNA fragments were pulled with the MyOne Streptavidin C1 beads (Life Technologies). To repair the ends of sheared DNA and remove biotin from unligated ends, DNA-bound beads were resuspended in 100µl of mix containing 82µl of 1X NEB T4 DNA ligase buffer with 10mM ATP (NEB), 10µl of 10 (2.5mM each) 25mM dNTP mix, 5µl of 10U/µl NEB T4 PNK (NEB), 4µl of 3U/µl NEB T4 DNA polymerase (NEB), and 1µl of 5U/µl NEB DNA polymerase I, Large (Klenow) Fragment (NEB).

After end-repair, dATP attachment was carried out in 100µl of mix consisting of 90µl of 1X NEBuffer 2, 5µl of 10mM dATP, 5µl of 5U/µl NEB Klenow exo minus (NEB), and incubated at 37C for 30 minutes. The beads were then cleaned for Illumina sequencing adaptor ligation which was done in a mix containing 50µl of 1X T4 ligase buffer, 3µl T4 DNA ligases (NEB), and 2µl of a 15µM Illumina indexed adapter at room temperature for 1 hour. DNA was dissociated from the bead by heating at 98C for 10 minutes, separated on a magnet, and transferred to a clean tube.

Final amplification of the library was carried out in multiple PCR reactions using Illumina PCR primers. The reactions were performed on a 25µl scale consisting of 25ng of DNA, 2µl of 2.5mM dNTPs, 0.35µl of 10µM each primer, 2.5µl of 10X PfuUltra buffer, and 0.5µl of PfuUltra II Fusion DNA polymerase (Agilent). The PCR cycle conditions were set to 98C for 30 seconds as the denaturing step, followed by 16 cycles of 98C 10 seconds, 65C for 30 seconds, 72C for 30 seconds, then with an extension step at 72C for 7 minutes.

After PCR amplification, the products from the same library were pooled and fragments ranging in sized of 300–500 bp were selected with AMPure XP beads (Beckman Coulter). The sized selected libraries were sequenced to produce paired-end Hi-C reads on the Illumina HiSeq 2500 platform with the V4 for 125 cycles.

### Hi-C Matrix Generation

Paired end reads were processed using the juicer pipeline with default parameters. Reads were mapped to reference genome hg19, with “-s” (site parameter) MboI. Reads with MAPQ >30 were kept for further analysis. Data was extracted and input to MATLAB using Juicebox tools command “dump”. Knight-Ruiz (KR) normalization was applied to all matrices, observed over expected (O/E) matrices were used for A/B compartmentalization and identification of topologically associating domains (TADs). Rows and columns for which more than 10% of entries had zeros were removed from the matrix.

### A/B Compartmentalization

Hi-C data was partitioned into A/B compartments according to the sign of the chromatin accessibility metric termed the Fiedler vector (the eigenvector corresponding to the second smallest eigenvalue of the normalized Laplacian matrix). Positive vector values were assigned to the ‘A’ compartment representing euchromatic loci and negative values were assigned to the ‘B’ compartment representing heterochromatic loci. The magnitude of the Fiedler values indicate the level of openness or closedness of the chromatin at a given loci. As chromatin evolves from one state of accessibility to another, it may switch compartments. Hi-C data was at 100kb resolution.

### TAD Calling

Topologically associating domains (TADs) were designated using spectral identification as described in [34]. Briefly, the Fiedler vector is calculated for a normalized Hi-C adjacency matrix and initial TADs are organized according to neighboring regions whose Fiedler values have the same sign. The initial TAD structure is repartitioned if for a given domain, the Fiedler number falls below a user-specified threshold (λthr). This ensures that TADs are not too large and repartitions the domains until the Fiedler number is larger than the threshold or until the TAD reaches the smallest allowable TAD size (default is 3). This process is performed iteratively for a set of chromosome-specific Hi-C contact maps. For our analysis, *λthr* was chosen to ensure a median TAD size of 900kb, as the expected median TAD size in mammalian genomes is 880kb (rounded to 900kb for our data since bins are in intervals of 100kb)[63]. *λthr* was chosen individually for each chromosome to ensure each TAD clustering set would have the same approximate median TAD size. Hi-C data was at 100kb resolution.

### Transcription Factor Enrichment Analysis

Transcription factor enrichment analysis was performed using oPOSSUM-3 software [37]. oPOSSUM-3 looks for the over-representation of transcription factor binding sites (TFBS) in the DNA sequences of a set of genes, using Z-score and Fisher exact probability to determine significance. For our analysis, 64 genes from a differentially conformed region of chromosome 19 were submitted as query and compared against a background set of genes in the oPOSSUM-3 database. Both sets of genes were compared to the JASPAR CORE database of preferential TFBS, representing 719 vertebrate TFBS, and the rate of TFBS occurrence for each set of genes was measured [64]. The following analysis parameters were used: 8 bit minimum profile specificity, 0.40 conservation cutoff, 85% matrix score threshold, and a 5kb flanking length.

### Synthetic 5C Map

We constructed a synthetic 5C contact map derived from a genome-wide 1Mb Hi-C contact map for genomic regions containing genes in the EMT and E2F gene networks. This was done by locating the genomic bins corresponding to the network genes, extracting the interchromosomal and network-specific intrachromosomal interaction frequencies for those bins, and stitching them together in genomic order. Our working set of network genes was sourced from the KEGG database and included 189 EMT genes and 189 E2F genes distributed across all chromosomes in the genome. Some of the 378 total genes occupy the same 1Mb genomic bins resulting in a 333 by 333 1Mb-resolution synthetic 5C adjacency matrix.

### Permutation Test

To determine whether the EMT and E2F gene networks are more closely interacting than other regions of the genome, we performed a permutation test with 1000 randomly sampled sets of 333 1Mb genomic bins across the genome. The Fiedler number for each set was computed and the distribution of numbers was plotted with the observed Fiedler number corresponding to the combined EMT and E2F gene networks. Our null hypothesis is that the EMT/E2F gene networks and random gene sets do not differ and our significance level is 0.1.

### Von Neumann Entropy

Entropy is a measure of “disorder” in a given system – the higher the entropy, the greater the disorder. In the context of genome structure, the higher the entropy, the more conformations available to the system. If the distant ends of a genomic region - say a single gene – interact to form a loop, there are fewer conformations available to the gene and thus the entropy of that genomic region is reduced. In this way, entropy can be considered a signature for chromatin conformation. We compute the Von Neumann Entropy (VNE) - multivariate entropy - using the following equation: 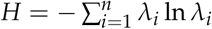, where *H* takes on positive values with larger values indicating higher chromatin accessibility.

## Figure and Table Legends

**Supplemental Figure 1.**
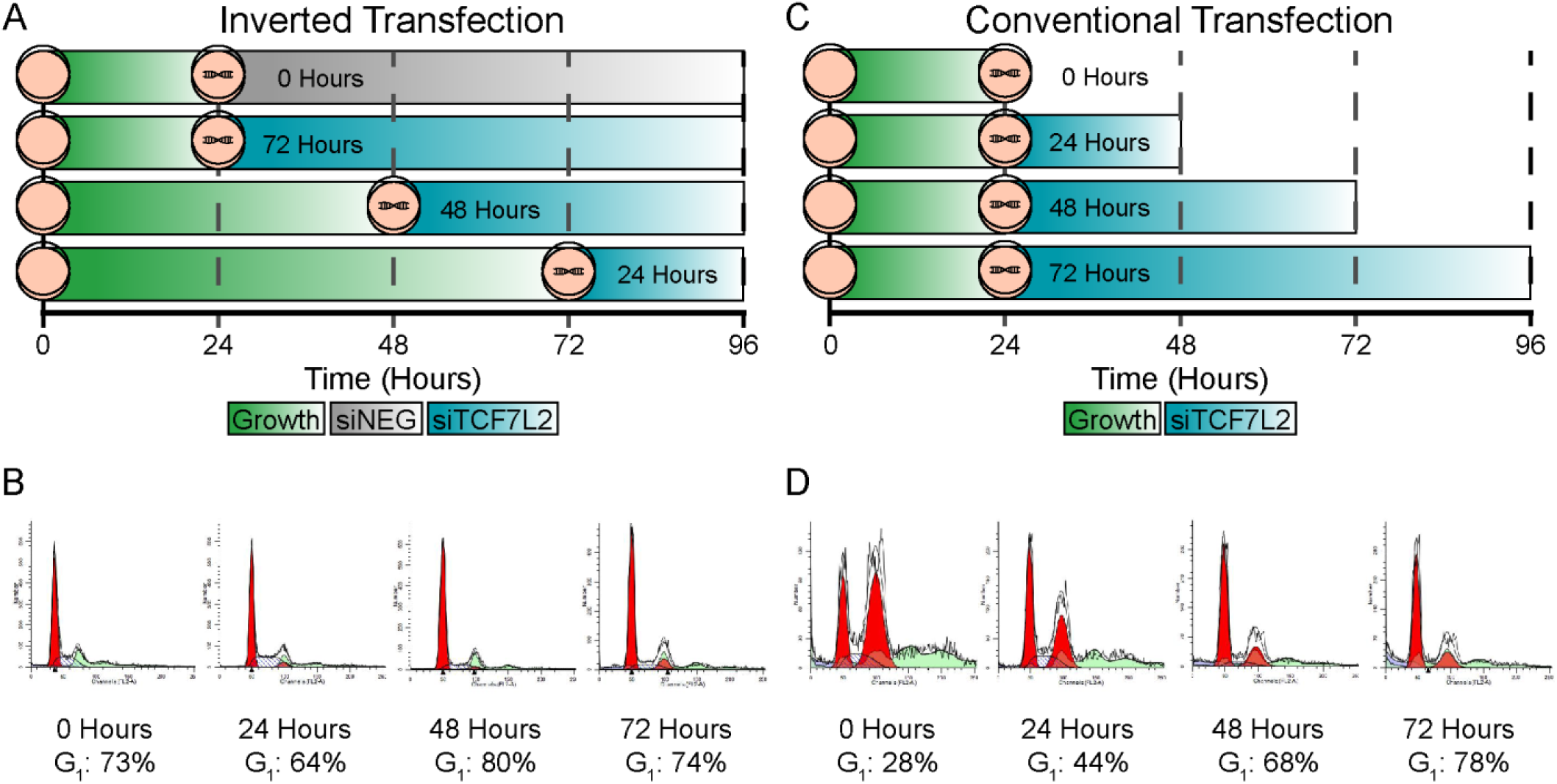
Validation of the Inverse Transfection Methodology. (A) Schematic demonstrating an Inverse Transfection methodology, in which the cells are grown equal amounts of time, with successive addition of siRNA at defined time points. Cells are then harvested simultaneously. The time course structure is generated by the amount of time incubated with siRNA. The time course was performed until 72 hours post-transfection based on the capability to synchronize cells by this time point, the silencing efficiency, as well as the results of a preliminary gene expression microarray, which demonstrated that the majority of changes in gene expression found at a later time point (96 hours post-transfection) first appeared at 72 hours post-transfection (results not shown). (B) The inverted transfection methodology resulted in ∼70% of the cells in the G_1_/G_0_ phase of the cell cycle across the time points, thereby permitting time series Hi-C analysis. The G_1_ is shown in red and is the leftmost peak on the graph. (C) Schematic demonstrating a conventional transfection methodology, which involves the simultaneous addition of siRNA to multiple cell culture vessels. The time course structure is generated by successive harvesting of the cells at defined time points from different vessels. (D) However, cells harvested at different time points are grown for different amounts of time, resulting in varying cell cycle distributions for each time point. The G_1_ peak is the leftmost peak in red, the G_2_ peak is the rightmost peak in red, which decreases in size as time progresses. S-phase is denoted by the angled lines against a clear background found between the G_1_ and G_2_ peaks. Cellular debris is shown in purple and aggregates in light green. Given that the number of cells in G_1_ varies from ∼20% at Time 0 to ∼80% at Time 72, we find this transfection methodology unsuitable for Hi-C analysis as differences in chromatin structure between the time points arise as a result of varying cell cycle distributions [**?**]. Note: When using a colorectal cancer cell line, such as SW480, or other cell line with a clumped growth pattern, the success of an Inverse Transfection is contingent upon the number of cells added to the vessel. If cells are added in excess, the transfection will suffer from low efficiency; conversely, if too few cells are added, the population will not undergo G_1_ arrest by contact inhibition. Fortuitously, *TCF7L2* silencing has been reported to result in a G_1_ arrest, supporting our aim [5]. Cells growing in a monolayer, such as RPE-1, can be added in excess without loss of transfection efficiency (data not shown.

**Supplemental Figure 2.**
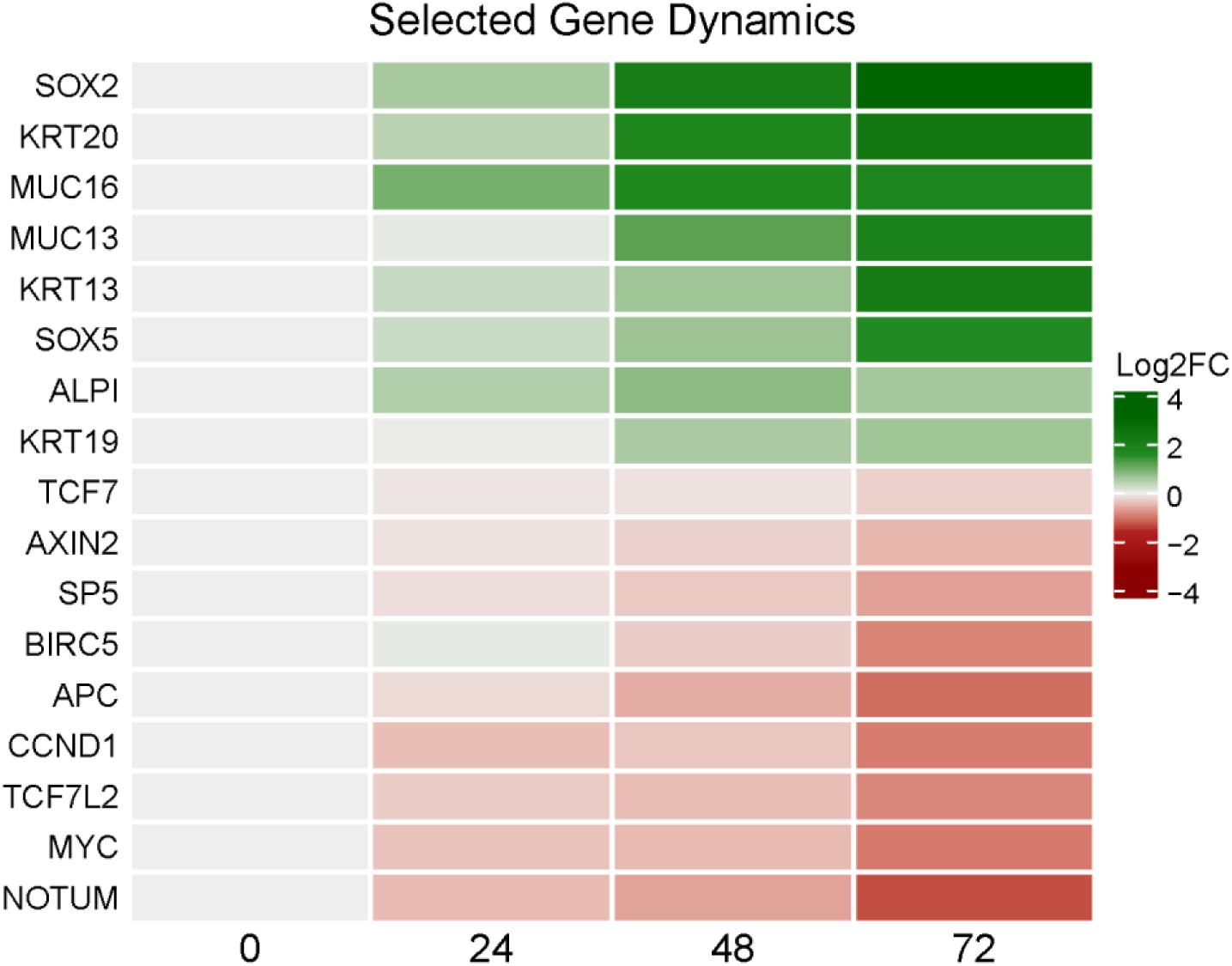
Influence of *TCF7L2* on Selected Genes of Interest Involved in Colorectal Cancer. To assess the influence of silencing *TCF7L2*, we examined the expression profiles of individual genes with known biological roles. Here, the log2FC in expression of each gene (compared to 0 hours) was plotted in a heatmap. Known target genes including *MYC* and *CCND1* show a similar expression profile to *TCF7L2*. Key negative regulators of Wnt signaling, *APC* and *NOTUM*, also show a similar pattern to *TCF7L2*, supporting a Wnt signaling intrinsic feedback loop. The loss of *NOTUM* may explain why the expression of *AXIN2* does not decrease to the same degree as *CCND1* or *MYC*. Known markers of intestinal differentiation, including genes from the mucin and keratin families, demonstrate increased expression upon silencing of *TCF7L2*.

**Supplemental Figure 3.**
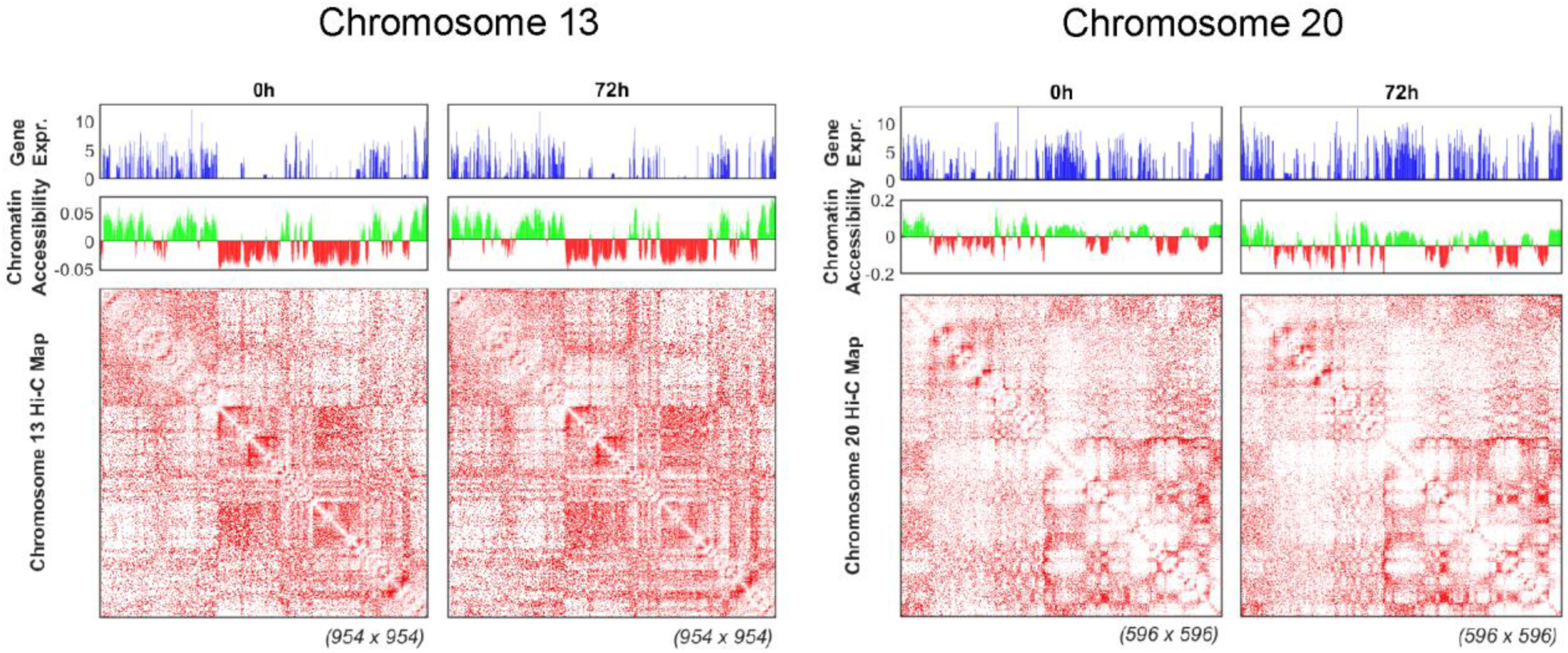
Interplay between Chromatin Structure and Gene Expression. The top panels show RNA expression (TPM) for chromosomes 13 and 20, while the middle panels show Fiedler vector computed from the OE normalized Hi-C matrices. Positive Fiedler values correspond to compartment A (euchromatin), whereas negative values correspond to compartment B (heterochromatin). The bottom panel consists of normalized Hi-C matrices. Each Hi-C contact map is a symmetric square matrix and the size of each map, denoted at the bottom right corner, refers to the number of 100kb genomic bins represented in the map (or equivalently, the number of rows and columns in the matrix). Overall, a close correlation between gene expression and chromatin conformation is observed.

**Supplemental Figure 4.**
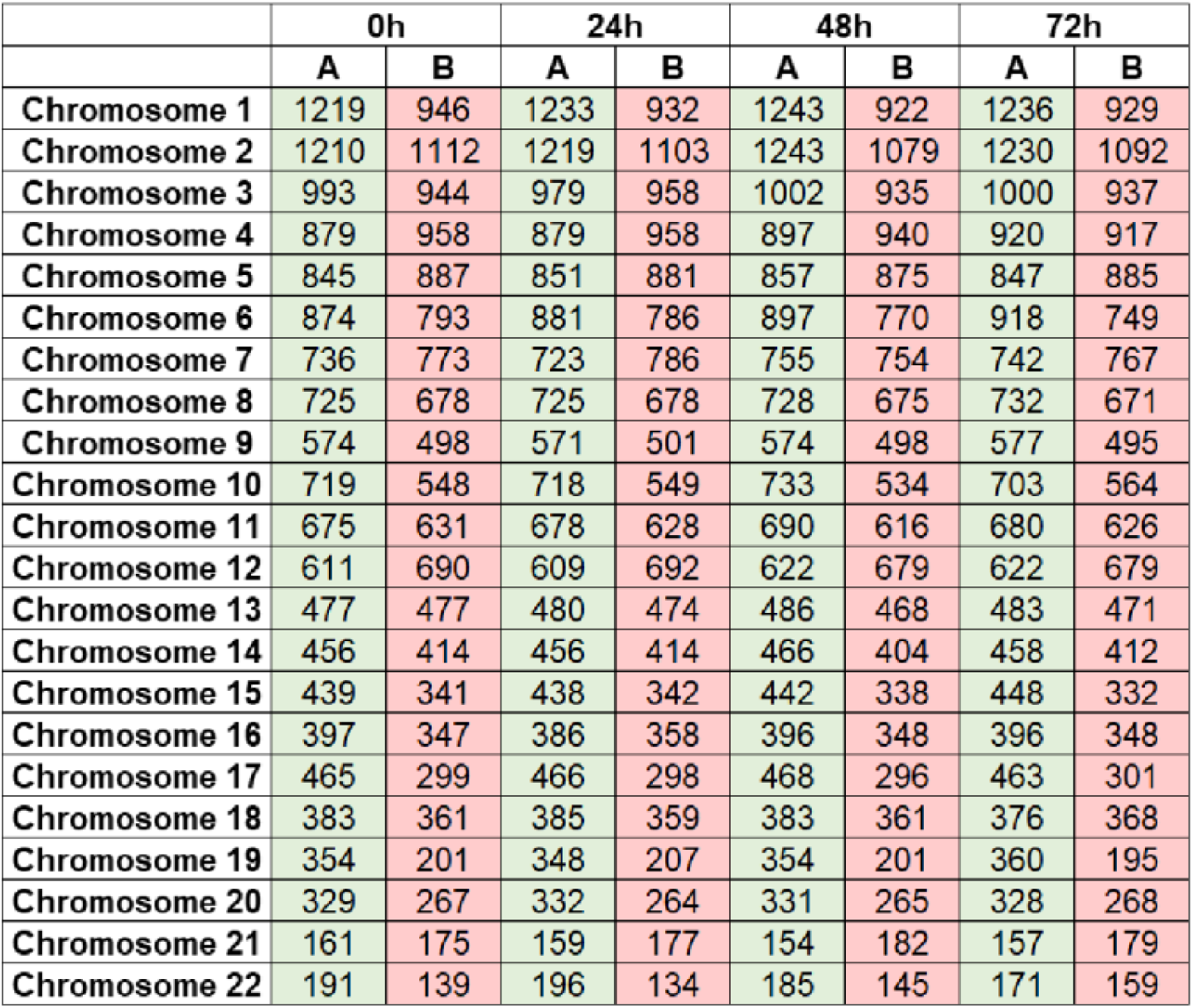
A/B Compartment Switching Over Time. Number of 100kb loci in “A” or “B” compartments at each time point for each chromosome. While ∼4% of the genome undergoes an A/B compartment switch at any time point, the ratio of A/B compartments remains the same.

**Supplemental Figure 5.**
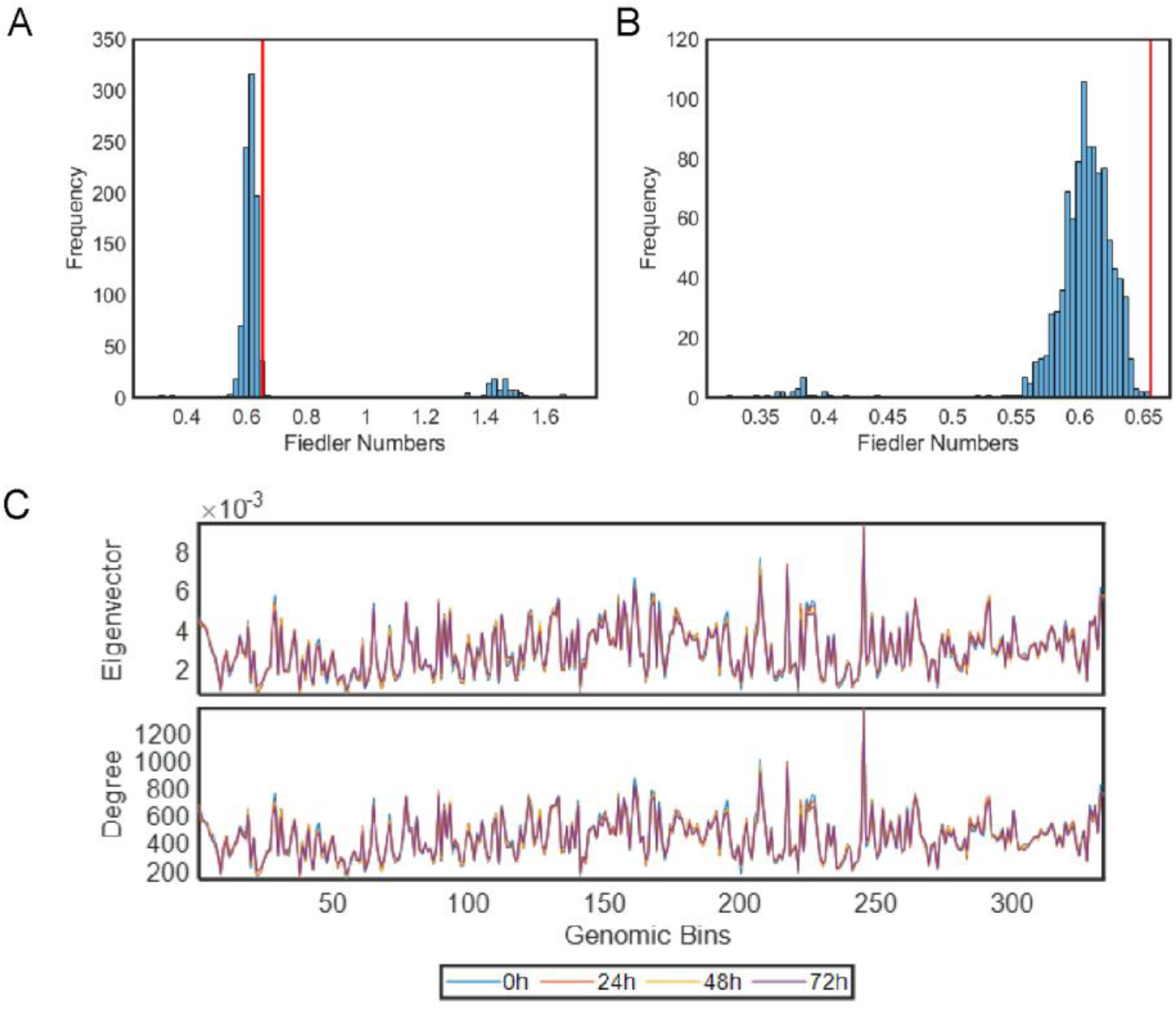
Permutation test distributions and Centrality Analyses. (A) Fiedler numbers (a measure of chromatin connectivity) for 1000 randomly sampled gene sets were generated and the distribution plotted (left). (B) Genes in the EMT and E2F networks have a significantly higher connectivity than genes in randomly sampled gene sets (*p* = 0.001)(right). (C) The Eigenvector and Degree centralities do not change significantly over time.

## References

[1] Vladimir Korinek, Nick Barker, Petra Moerer, Elly van Donselaar, Gerwin Huls, Peter J Peters, and Hans Clevers. Depletion of epithelial stem-cell compartments in the small intestine of mice lacking tcf-4. Nature genetics, 19(4):379–383, 1998.

[2] Johan H van Es, Andrea Haegebarth, Pekka Kujala, Shalev Itzkovitz, Bon-Kyoung Koo, Sylvia F Boj, Jeroen Korving, Maaike van den Born, Alexander van Oudenaarden, Sylvie Robine, et al. A critical role for the wnt effector tcf4 in adult intestinal homeostatic selfrenewal. Molecular and cellular biology, 32(10):1918–1927, 2012.

[3] Nick Barker, Johan H Van Es, Jeroen Kuipers, Pekka Kujala, Maaike Van Den Born, Miranda Cozijnsen, Andrea Haegebarth, Jeroen Korving, Harry Begthel, Peter J Peters, et al. Identification of stem cells in small intestine and colon by marker gene lgr5. Nature, 449(7165):1003–1007, 2007.

[4] Toshiro Sato, Johan H Van Es, Hugo J Snippert, Daniel E Stange, Robert G Vries, Maaike Van Den Born, Nick Barker, Noah F Shroyer, Marc Van De Wetering, and Hans Clevers. Paneth cells constitute the niche for lgr5 stem cells in intestinal crypts. Nature, 469(7330):415–418, 2011.

[5] Marc Van De Wetering, Elena Sancho, Cornelis Verweij, Wim De Lau, Irma Oving, Adam Hurlstone, Karin Van Der Horn, Eduard Batlle, Damien Coudreuse, Anna-Pavlina Haramis, et al. The *β*-catenin/tcf-4 complex imposes a crypt progenitor phenotype on colorectal cancer cells. Cell, 111(2):241–250, 2002.

[6] Theodora Agalioti, Stavros Lomvardas, Bhavin Parekh, Junming Yie, Tom Maniatis, and Dimitris Thanos. Ordered recruitment of chromatin modifying and general transcription factors to the ifn-*β* promoter. Cell, 103(4):667–678, 2000.

[7] Andreas Hecht, Kris Vleminckx, Marc P Stemmler, Frans Van Roy, and Rolf Kemler. The p300/cbp acetyltransferases function as transcriptional coactivators of *β*-catenin in vertebrates. The EMBO journal, 19(8):1839–1850, 2000.

[8] Nick Barker, Adam Hurlstone, Hannah Musisi, Antony Miles, Mariann Bienz, and Hans Clevers. The chromatin remodelling factor brg-1 interacts with *β*-catenin to promote target gene activation. The EMBO journal, 20(17):4935–4943, 2001.

[9] Christian Mosimann, George Hausmann, and Konrad Basler. Parafibromin/hyrax activates wnt/wg target gene transcription by direct association with *β*-catenin/armadillo. Cell, 125(2):327–341, 2006.

[10] Kenneth W Kinzler and Bert Vogelstein. Lessons from hereditary colorectal cancer. Cell, 87(2):159–170, 1996.

[11] AJ Rowan, H Lamlum, M Ilyas, J Wheeler, J Straub, A Papadopoulou, D Bicknell, WF Bodmer, and IPM Tomlinson. Apc mutations in sporadic colorectal tumors: a mutational “hotspot” and interdependence of the “two hits”. Proceedings of the National Academy of Sciences, 97(7):3352–3357, 2000.

[12] Patrice J Morin, Andrew B Sparks, Vladimir Korinek, Nick Barker, Hans Clevers, Bert Vogelstein, and Kenneth W Kinzler. Activation of *β*-catenin-tcf signaling in colon cancer by mutations in *β*-catenin or apc. Science, 275(5307):1787–1790, 1997.

[13] Vladimir Korinek, Nick Barker, Patrice J Morin, Dick Van Wichen, Roel De Weger, Kenneth W Kinzler, Bert Vogelstein, and Hans Clevers. Constitutive transcriptional activation by a *β*-catenin-tcf complex in apc-/-colon carcinoma. Science, 275(5307):1784–1787, 1997.

[14] Tong-Chuan He, Andrew B Sparks, Carlo Rago, Heiko Hermeking, Leigh Zawel, Luis T Da Costa, Patrice J Morin, Bert Vogelstein, and Kenneth W Kinzler. Identification of c-myc as a target of the apc pathway. Science, 281(5382):1509–1512, 1998.

[15] Osamu Tetsu and Frank McCormick. *β*-catenin regulates expression of cyclin d1 in colon carcinoma cells. Nature, 398(6726):422–426, 1999.

[16] David S Parker, Yunyun Y Ni, Jinhee L Chang, Jiong Li, and Ken M Cadigan. Wingless signaling induces widespread chromatin remodeling of target loci. Molecular and cellular biology, 28(5):1815–1828, 2008.

[17] Erez Lieberman-Aiden, Nynke L Van Berkum, Louise Williams, Maxim Imakaev, Tobias Ragoczy, Agnes Telling, Ido Amit, Bryan R Lajoie, Peter J Sabo, Michael O Dorschner, et al. Comprehensive mapping of long-range interactions reveals folding principles of the human genome. science, 326(5950):289–293, 2009.

[18] Jesse R Dixon, Siddarth Selvaraj, Feng Yue, Audrey Kim, Yan Li, Yin Shen, Ming Hu, Jun S Liu, and Bing Ren. Topological domains in mammalian genomes identified by analysis of chromatin interactions. Nature, 485(7398):376–380, 2012.

[19] Suhas SP Rao, Miriam H Huntley, Neva C Durand, Elena K Stamenova, Ivan D Bochkov, James T Robinson, Adrian L Sanborn, Ido Machol, Arina D Omer, Eric S Lander, et al. A 3d map of the human genome at kilobase resolution reveals principles of chromatin looping. Cell, 159(7):1665–1680, 2014.

[20] Robert-Jan Palstra, Bas Tolhuis, Erik Splinter, Rian Nijmeijer, Frank Grosveld, and Wouter de Laat. The *β*-globin nuclear compartment in development and erythroid differentiation. Nature genetics, 35(2):190–194, 2003.

[21] Indika Rajapakse and Mark Groudine. On emerging nuclear order. Journal of Cell Biology, 192(5):711–721, 2011.

[22] Thomas Ried and Indika Rajapakse. The 4d nucleome. Methods (San Diego, Calif.), 123:1, 2017.

[23] Scott Ronquist, Geoff Patterson, Lindsey A Muir, Stephen Lindsly, Haiming Chen, Markus Brown, Max S Wicha, Anthony Bloch, Roger Brockett, and Indika Rajapakse. Algorithm for cellular reprogramming. Proceedings of the National Academy of Sciences, 114(45):11832–11837, 2017.

[24] Harold Weintraub, Stephen J Tapscott, Robert L Davis, Mathew J Thayer, Mohammed A Adam, Andrew B Lassar, and A Dusty Miller. Activation of muscle-specific genes in pigment, nerve, fat, liver, and fibroblast cell lines by forced expression of myod. Proceedings of the National Academy of Sciences, 86(14):5434–5438, 1989.

[25] Kazutoshi Takahashi and Shinya Yamanaka. Induction of pluripotent stem cells from mouse embryonic and adult fibroblast cultures by defined factors. cell, 126(4):663–676, 2006.

[26] Kazutoshi Takahashi, Koji Tanabe, Mari Ohnuki, Megumi Narita, Tomoko Ichisaka, Kiichiro Tomoda, and Shinya Yamanaka. Induction of pluripotent stem cells from adult human fibroblasts by defined factors. cell, 131(5):861–872, 2007.

[27] B Michael Ghadimi, Marian Grade, Michael J Difilippantonio, Sudhir Varma, Richard Simon, Cristina Montagna, Laszlo Füzesi, Claus Langer, Heinz Becker, Torsten Liersch, et al. Effectiveness of gene expression profiling for response prediction of rectal adenocarcinomas to preoperative chemoradiotherapy. Journal of clinical oncology: official journal of the American Society of Clinical Oncology, 23(9):1826, 2005.

[28] Emil Kendziorra, Kerstin Ahlborn, Melanie Spitzner, Margret Rave-Fränk, Georg Emons, Jochen Gaedcke, Frank Kramer, Hendrik A Wolff, Heinz Becker, Tim Beissbarth, et al. Silencing of the wnt transcription factor tcf4 sensitizes colorectal cancer cells to (chemo-) radiotherapy. Carcinogenesis, 32(12):1824–1831, 2011.

[29] D Cunningham. atkin w, lenz h, lynch ht, minsky b, nordlinger b, et al. colorectal cancer. The Lancet, 375(9719):1030–1047, 2010.

[30] Gilbert Weidinger, Chris J Thorpe, Katrin Wuennenberg-Stapleton, John Ngai, and Randall T Moon. The sp1-related transcription factors sp5 and sp5-like act downstream of wnt/*β*-catenin signaling in mesoderm and neuroectoderm patterning. Current biology, 15(6):489–500, 2005.

[31] Meiko Takahashi, Yusuke Nakamura, Kazutaka Obama, and Yoichi Furukawa. Identification of sp5 as a downstream gene of the *β*-catenin/tcf pathway and its enhanced expression in human colon cancer. International journal of oncology, 27(6):1483–1487, 2005.

[32] Pantelis Hatzis, Laurens G van der Flier, Marc A van Driel, Victor Guryev, Fiona Nielsen, Sergei Denissov, Isaäc J Nijman, Jan Koster, Evan E Santo, Willem Welboren, et al. Genome-wide pattern of tcf7l2/tcf4 chromatin occupancy in colorectal cancer cells. Molecular and cellular biology, 28(8):2732–2744, 2008.

[33] Yusuke Kamachi, Masanori Uchikawa, and Hisato Kondoh. Pairing sox off: with partners in the regulation of embryonic development. Trends in Genetics, 16(4):182–187, 2000.

[34] Jie Chen, Alfred O Hero III, and Indika Rajapakse. Spectral identification of topological domains. Bioinformatics, 32(14):2151–2158, 2016.

[35] Peter R Cook and Davide Marenduzzo. Transcription-driven genome organization: a model for chromosome structure and the regulation of gene expression tested through simulations. Nucleic acids research, 46(19):9895–9906, 2018.

[36] Nicole Beauchemin and Azadeh Arabzadeh. Carcinoembryonic antigen-related cell adhesion molecules (ceacams) in cancer progression and metastasis. Cancer and Metastasis Reviews, 32(3-4):643–671, 2013.

[37] Andrew T Kwon, David J Arenillas, Rebecca Worsley Hunt, and Wyeth W Wasserman. opossum-3: advanced analysis of regulatory motif over-representation across genes or chip-seq datasets. G3: Genes, Genomes, Genetics, 2(9):987–1002, 2012.

[38] Aravind Subramanian, Pablo Tamayo, Vamsi K Mootha, Sayan Mukherjee, Benjamin L Ebert, Michael A Gillette, Amanda Paulovich, Scott L Pomeroy, Todd R Golub, Eric S Lander, et al. Gene set enrichment analysis: a knowledge-based approach for interpreting genome-wide expression profiles. Proceedings of the National Academy of Sciences, 102(43):15545–15550, 2005.

[39] Alexey Sergushichev. An algorithm for fast preranked gene set enrichment analysis using cumulative statistic calculation. BioRxiv, page 060012, 2016.

[40] Seiichi Ishida, Erich Huang, Harry Zuzan, Rainer Spang, Gustavo Leone, Mike West, and Joseph R Nevins. Role for e2f in control of both dna replication and mitotic functions as revealed from dna microarray analysis. Molecular and cellular biology, 21(14):4684–4699, 2001.

[41] Bing Ren, Hieu Cam, Yasuhiko Takahashi, Thomas Volkert, Jolyon Terragni, Richard A Young, and Brian David Dynlacht. E2f integrates cell cycle progression with dna repair, replication, and g2/m checkpoints. Genes & development, 16(2):245–256, 2002.

[42] Steven M Chan, Andrew P Weng, Robert Tibshirani, Jon C Aster, and Paul J Utz. Notch signals positively regulate activity of the mtor pathway in t-cell acute lymphoblastic leukemia. Blood, The Journal of the American Society of Hematology, 110(1):278–286, 2007.

[43] Jing Xu, Yongjun Dang, Yunzhao R Ren, and Jun O Liu. Cholesterol trafficking is required for mtor activation in endothelial cells. Proceedings of the National Academy of Sciences, 107(10):4764–4769, 2010.

[44] Anne Boulay, Joelle Rudloff, Jingjing Ye, Sabine Zumstein-Mecker, Terence O’Reilly, Dean B Evans, Shiuan Chen, and Heidi A Lane. Dual inhibition of mtor and estrogen receptor signaling in vitro induces cell death in models of breast cancer. Clinical Cancer Research, 11(14):5319–5328, 2005.

[45] Gilbert Strang, Gilbert Strang, Gilbert Strang, and Gilbert Strang. Introduction to linear algebra, volume 3. Wellesley-Cambridge Press Wellesley, MA, 1993.

[46] Laura Seaman, Haiming Chen, Markus Brown, Darawalee Wangsa, Geoff Patterson, Jordi Camps, Gilbert S Omenn, Thomas Ried, and Indika Rajapakse. Nucleome analysis reveals structure–function relationships for colon cancer. Molecular Cancer Research, 15(7):821–830, 2017.

[47] Sijia Liu, Haiming Chen, Scott Ronquist, Laura Seaman, Nicholas Ceglia, Walter Meixner, Pin-Yu Chen, Gerald Higgins, Pierre Baldi, Steve Smale, et al. Genome architecture mediates transcriptional control of human myogenic reprogramming. iScience, 6:232–246, 2018.

[48] Sijia Liu, Pin-Yu Chent, Indika Rajapakse, and Alfred Hero. First-order bifurcation detection for dynamic complex networks. In 2018 IEEE International Conference on Acoustics, Speech and Signal Processing (ICASSP), pages 6912–6916. IEEE, 2018.

[49] Pin-Yu Chen, Lingfei Wu, Sijia Liu, and Indika Rajapakse. Fast incremental von neumann graph entropy computation: Theory, algorithm, and applications. arXiv preprint arXiv:1805.11769, 2018.

[50] Stephen P Borgatti. Centrality and network flow. Social networks, 27(1):55–71, 2005.

[51] MEJ Newman. Networks: An introduction. 2010 oxford.

[52] Abdolrahman S Nateri, Bradley Spencer-Dene, and Axel Behrens. Interaction of phosphorylated c-jun with tcf4 regulates intestinal cancer development. Nature, 437(7056):281–285, 2005.

[53] Tomas Vacik, Jennifer L Stubbs, and Greg Lemke. A novel mechanism for the transcriptional regulation of wnt signaling in development. Genes & development, 25(17):1783–1795, 2011.

[54] Ken M Cadigan and Marian L Waterman. Tcf/lefs and wnt signaling in the nucleus. Cold Spring Harbor perspectives in biology, 4(11):a007906, 2012.

[55] Shuichi Hashimoto, Huaiyong Chen, Jianwen Que, Brian L Brockway, Jeffrey A Drake, Joshua C Snyder, Scott H Randell, and Barry R Stripp. *β*-catenin–sox2 signaling regulates the fate of developing airway epithelium. Journal of cell science, 125(4):932–942, 2012.

[56] Abby Sarkar, Aaron J Huebner, Rita Sulahian, Anthony Anselmo, Xinsen Xu, Kyle Flattery, Niyati Desai, Carlos Sebastian, Mary Anna Yram, Katrin Arnold, et al. Sox2 suppresses gastric tumorigenesis in mice. Cell reports, 16(7):1929–1941, 2016.

[57] Ralph Stadhouders, Enrique Vidal, François Serra, Bruno Di Stefano, François Le Dily, Javier Quilez, Antonio Gomez, Samuel Collombet, Clara Berenguer, Yasmina Cuartero, et al. Transcription factors orchestrate dynamic interplay between genome topology and gene regulation during cell reprogramming. Nature genetics, 50(2):238–249, 2018.

[58] Rüdiger Braun, Lena Anthuber, Daniela Hirsch, Darawalee Wangsa, Justin Lack, Nicole E McNeil, Kerstin Heselmeyer-Haddad, Irianna Torres, Danny Wangsa, Markus A Brown, et al. Single cell-derived primary rectal carcinoma cell lines reflect intratumor heterogeneity associated with treatment response. Clinical Cancer Research, 2020.

[59] Francis Blokzijl, Joep De Ligt, Myrthe Jager, Valentina Sasselli, Sophie Roerink, Nobuo Sasaki, Meritxell Huch, Sander Boymans, Ewart Kuijk, Pjotr Prins, et al. Tissuespecific mutation accumulation in human adult stem cells during life. Nature, 538(7624):260–264, 2016.

[60] Helmuth Gehart and Hans Clevers. Tales from the crypt: new insights into intestinal stem cells. Nature Reviews Gastroenterology & Hepatology, 16(1):19–34, 2019.

[61] B Mann, M Gelos, A Siedow, ML Hanski, A Gratchev, M Ilyas, WF Bodmer, MP Moyer, EO Riecken, HJ Buhr, et al. Target genes of *β*-catenin–t cell-factor/lymphoidenhancer-factor signaling in human colorectal carcinomas. Proceedings of the National Academy of Sciences, 96(4):1603–1608, 1999.

[62] Xiao-qing Gan, Ji-yong Wang, Ying Xi, Zhi-li Wu, Yiping Li, and Lin Li. Nuclear dvl, c-jun, *β*-catenin, and tcf form a complex leading to stabilization of *β*-catenin–tcf interaction. The Journal of cell biology, 180(6):1087–1100, 2008.

[63] Boyan Bonev and Giacomo Cavalli. Organization and function of the 3d genome. Nature Reviews Genetics, 17(11):661, 2016.

[64] Aziz Khan, Oriol Fornes, Arnaud Stigliani, Marius Gheorghe, Jaime A Castro-Mondragon, Robin van der Lee, Adrien Bessy, Jeanne Cheneby, Shubhada R Kulkarni, Ge Tan, et al. Jaspar 2018: update of the open-access database of transcription factor binding profiles and its web framework. Nucleic acids research, 46(D1):D260–D266, 2018.

